# The impact of insecticide decay on the rate of insecticide resistance evolution for monotherapies and mixtures

**DOI:** 10.1101/2024.05.03.592309

**Authors:** Neil Philip Hobbs, Ian Michael Hastings

## Abstract

The issue of insecticide decay in the public health deployments of insecticides is frequently highlighted as an issue for disease control. There are additional concerns insecticide decay also impacts the selection for insecticide resistance. Despite these concerns insecticide decay is lacking from models evaluating insecticide resistance management strategies. The impact of insecticide decay is modelled using a model which assumes a polygenic basis of insecticide resistance. Single generation selection events covering the insecticide efficacy and insecticide resistance space for both monotherapies and mixtures are conducted. With the outcome being the between generation change in the bioassay survival to the insecticides. The monotherapy sequence strategy and mixture strategy were compared against each other when including insecticide decay, with the outcome being the difference in strategy lifespan. The results demonstrate that as insecticides decay, they can apply a greater selection pressure than newly deployed insecticides, a process which can occur for both monotherapies and mixtures. For mixtures, it is seen that the rate of selection is highest when both insecticides are at reduced efficacies which would occur if reduced dose mixtures were used. Inclusion of insecticide decay in simulations was found to reduce the benefit of mixtures against monotherapy sequences, and this is especially so when reduced-dose mixtures are used. Insecticide decay is often highlighted as an important consideration for mixtures. The inclusion of insecticide decay in models is often lacking, and these results indicate this is absence may be over-inflating the performance of full-dose mixtures. As insecticides decay, they still provide selection pressures with reduced ability to control transmission, replenishing worn-out insecticides more frequently should be considered.

## Introduction

Insecticides used for controlling malaria and other vector borne diseases often have long residual lifespans. For example, long-lasting insecticide-treated nets (LLINs) are expected to have an effective longevity of three years, while indoor residual sprays (IRS) are expected to have a lifespan between 3 and 9 months (WHO, 2012). The long durations between deployments means there is ample time for insecticides to decay and reduce in efficacy.

Insecticide decay is a concern for vector control because as insecticides decay over the months and years post-deployment, they will inevitably become less effective at killing the target vector population. Insecticide decay is frequently highlighted as an issue for disease control programmes, and as an effect that may undermine insecticide resistance management (IRM) strategies (Sternberg & Thomas, 2018). In contrast, decaying insecticide effectiveness is not regarded as an important issue in agricultural applications because insecticides rapidly degrade or are washed off crops by rain, both of which are generally regarded as beneficial in meeting consumer pressure for insecticide residues not being present on food.

Mathematical models are frequently used for evaluating IRM strategies in both agriculture and public health (Rex Consortium, 2007) and insecticide decay is generally absent (South et al., 2020), especially for the evaluation of strategies in a public health context where insecticide decay is considered an important issue. A systematic review of the resistance management literature (Rex Consortium, 2013) highlighted that when two insecticides are deployed in mixture they should have similar residual efficacies, however neither of the studies supporting this statement explicitly explored this (Curtis & Hill, 1993; Roush, 1989). Instead, this statement has often been considered an intuitive condition for the use of mixtures.

The important questions concern how insecticide decay influences the rate of evolution of insecticide resistance, and subsequently the effectiveness of IRM strategies? The implication of insecticide decay for IRM was considered by South et al. (2020) who highlighted there can be “windows of selection” during a deployment. This was considering resistance as a monogenic trait (i.e., RR, RS, SS) and considered only monotherapy deployments. Where insecticide decay is additionally highlighted as a main issue is mixtures (Curtis, 1985) because mixtures may become counter-productive as effectiveness of the constituent insecticides starts to decline. This implies that a mixture may be highly effective when first deployed, but counter-productive after a few months when effectiveness has fallen. Given the current trend for mixture LLINs, this needs to be addressed as a matter of some urgency.

Insecticide mixtures are often identified as effective IRM strategies for the public health deployment of insecticides (e.g., Curtis, 1985; Hobbs et al., 2023; Madgwick & Kanitz, 2022b; Mani, 1985; South & Hastings, 2018) when compared against monotherapy deployments. Although simulations suggest mixtures are only effective for IRM when both constituent insecticides are fully effective (South & Hastings, 2018); the general explanation for this being both need to be effective to provide mutual protection to the other (e.g., if an insect is resistant to insecticide *i*, then it needs to be reliably killed by insecticide *j* in the mixture).

Using a model which assumes a polygenic basis for insecticide resistance, a conceptual look at the implication of insecticide decay for both monotherapy and mixture deployments is taken. The aim of this paper is to understand and conceptualise the impact of insecticide decay on the IRM ability of monotherapies and mixtures when considering polygenic resistance and consider how the inclusion of insecticide decay in simulations may alter the preferable choice of IRM strategy.

## Methods

### Model Overview

A quantitative genetics model is used as previously described (Hobbs & Hastings, 2024) which assumes insecticide resistance is a polygenic trait. The level of insecticide resistance is quantified by the “polygenic resistance score” (PRS, *Z*_I_) which is measurable in standardised bioassays such as WHO cylinders. In the model, insecticide selection can be implemented by either a truncating or probabilistic selection process. Here the truncation selection branch of the model (“polytruncate”) is used as this branch allows for a better mechanistic understanding of the selection process with respect to insecticide decay. The key equation for interpretation of this paper is a modified sex-specific Breeder’s equation (Equation 3c in Hobbs & Hastings, 2024):

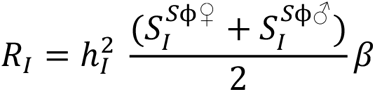

Where *R*_I_ is the response to selection and is the between generation change in the mean “polygenic resistance score” of the population. The values *S_I_*^Sϕ^*^♀^* and *S_I_*^Sϕ^*^♂^* are the selection differentials for females and males when considering resistance to insecticide *i*. Lowercase letters (*i*, *j*) indicate the insecticide and the uppercase letters (*I*, *J*) the corresponding resistance trait. The Breeder’s Equation is performed separately also for each insecticide and the corresponding resistance trait. The selection differentials (*S_I_*^Sϕ^*^♀^* and *S_I_*^Sϕ^*^♂^*) result from insecticide exposure (ϕ superscript) and fitness costs (ϕ superscript). The selection differential is the within generation change in the mean PRS, i.e., the difference in mean PRS between the newborns in the generation (*Z̄*_I_), and those individuals who survive to become parents of the next generation (*Z̄_I_*^P^). As male and female mosquitoes likely have different exposures to insecticides, the selection differentials are calculated separately (denoted by the superscripts *♀* for females and *♂* for males). ℎ*_I_*^2^ is the heritability of the trait. The value β is the exposure scaling factor and is used to calibrate simulations to expected timescales to account for uncertainty in values of heritability and the selection differentials. Figure 1 illustrates how insecticide deployment determines the selection differential.

**Figure 1:**
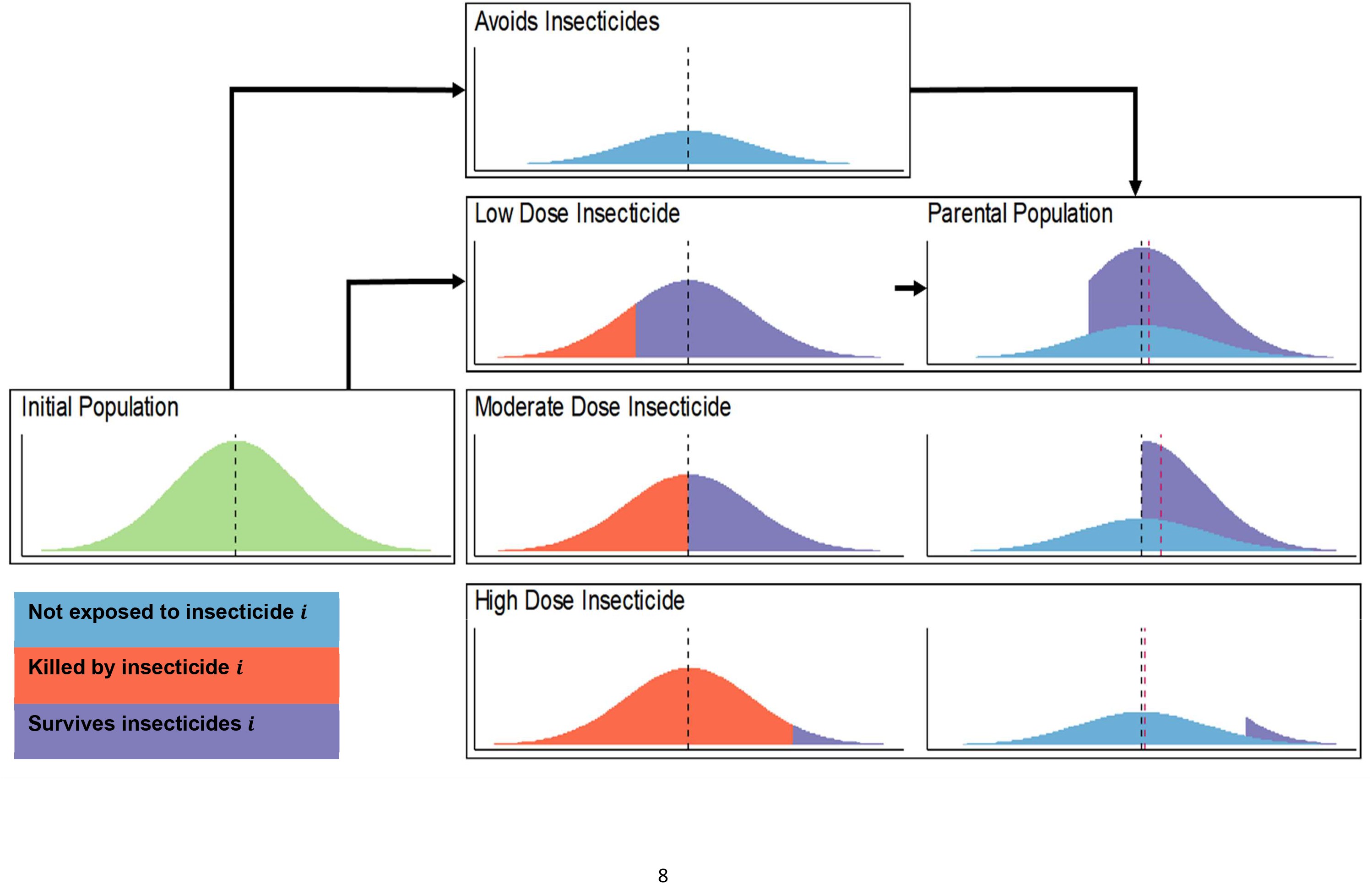
Mechanistic Explanation of the Impact of Insecticide Dose on the Rate of Selection. For all plots the x axis is the value of the “polygenic resistance score” (*Z*_I_) and the y axis is its frequency distribution in the population. The mosquito population initially emerges with a mean “polygenic resistance score” of *Z̄*_I_ (black dashed line). A proportion of these mosquitoes will avoid the insecticide(s) and have no selection pressure applied to them. The other proportion of the population will be exposed to the insecticide. The number surviving will depend on both the dose of insecticide mosquitoes are exposed to (such that higher insecticide doses kill more mosquitoes), and the level of resistance in the population (where higher resistance populations can better survive insecticides). The selection differential is the change in the mean “polygenic resistance score” between the initial emerging population (black dashed line, *Z̄*_I_) and the final parental population (red dashed line, *Z̄*_I_^P^), which is used in the Breeder’s Equation. In scenarios where most mosquitoes survive the insecticide, (e.g., the low dose example) the selection differential is low because many of the less resistant mosquitoes survive and is still further diluted by those mosquitoes escaping insecticide exposure. In scenarios where few mosquitoes survive the insecticide, (e.g., the high dose example) the selection differential is also low. This is because while those which survive the insecticide exposure are highly resistant, there are very few of these individuals. The unexposed individuals therefore make up the majority of the final parental population sufficiently diluting the highly resistant survivors. In the intermediary scenarios (e.g., the moderate dose example), the insecticide supplies sufficient selection pressure such that many moderately resistant mosquitoes survive. The number of mosquitoes surviving the insecticide is also sufficiently large such that the parental population is not sufficiently diluted by mosquitoes which avoided the insecticide(s).

The model is used to explore insecticide decay in two ways. First selection is implemented only on a single generation as this allows for a detailed understanding of the direct implication of insecticide decay allowing a detailed mechanistic understanding of the underlying selection process resulting from insecticide decay (Figures 1 and 2). Second simplistic, multi-generational scenarios of IRM strategy deployments are used to explore how including insecticide decay over longer timescales impacts the comparative ability of monotherapies and mixtures to delay the spread of IR over a 50 year time horizon.

**Figure 2:**
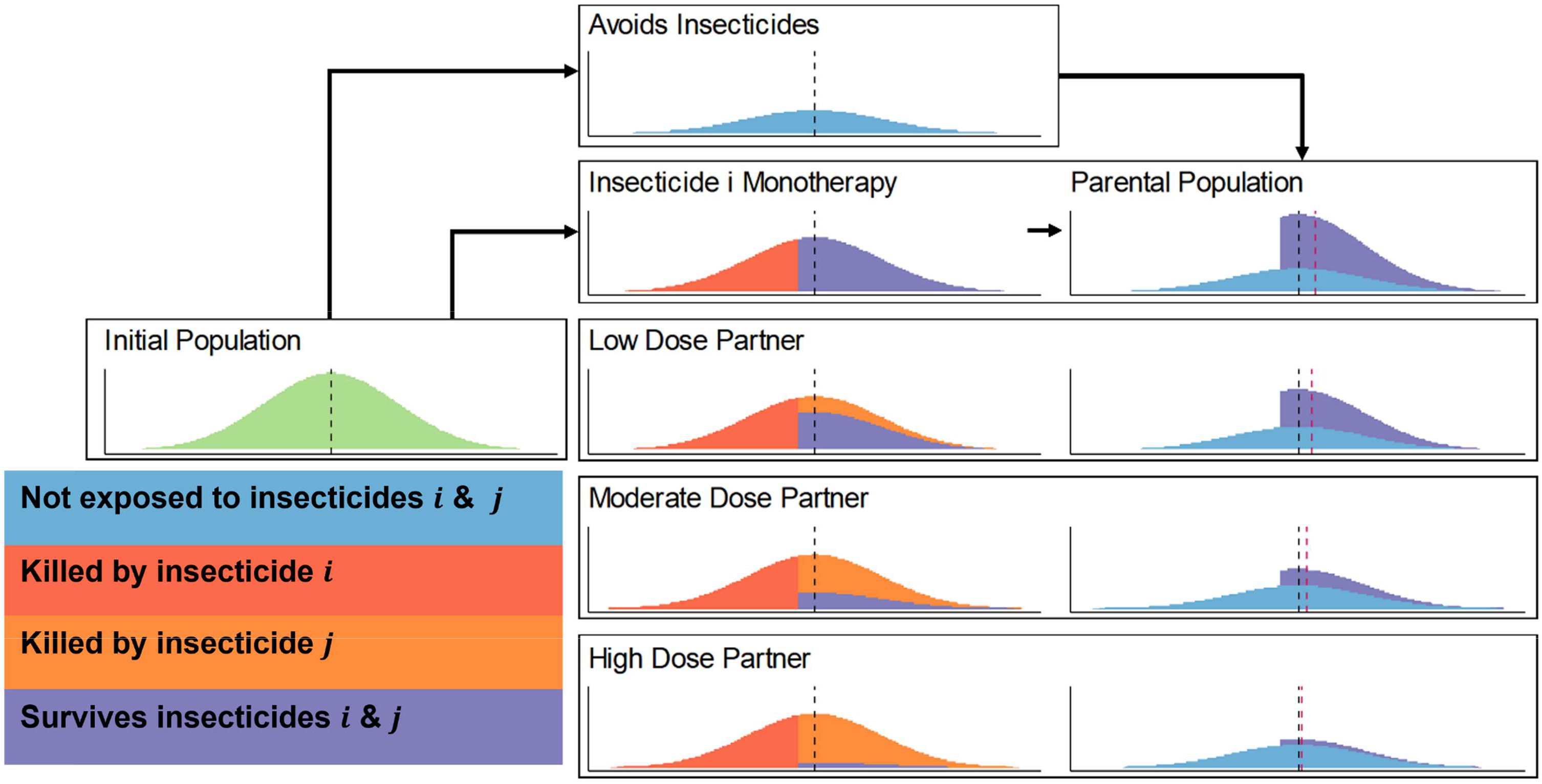
Mechanistic Explanation of the Impact of Insecticide Dose on the Rate of Selection: Mixtures. For all plots the x axis is the value of the “polygenic resistance score” (*Z*_I_) and the y axis is the frequency in the population. The initial mosquito population emerges with a mean resistance of *Z̄*_I_ to insecticide *i* and a mean resistance of *Z̄*_J_ to insecticide *j*. A proportion of these mosquitoes will avoid the insecticide(s) and have no selection pressure applied to them. The other proportion of the population will be exposed to the insecticide(s). The number surviving will depend on the dose of insecticide (such that higher insecticide doses kill more mosquitoes), and the level of resistance in the population (higher resistance populations can better survive insecticides). For scenarios where mosquitoes are exposed to a mixture insecticide *i* kills a proportion of those mosquitoes, depending on the level of resistance to insecticide *i* and the dosage of insecticide *i* (red). Those which survive insecticide *i* then also must survive insecticide *j* (which depends on the level of resistance to insecticide *j* and the dosage of insecticide *j*), and therefore insecticide *j* kills an additional proportion of the mosquitoes (orange), and therefore to become parents, mosquitoes most survive both insecticides (purple). We can then use this conceptual framework to explain how mixtures perform. If the partner insecticide (*j*) is not effective (low dose and/or high levels of resistance) then only a small additional number of mosquitoes are killed, not reducing the selection differential significantly compared to insecticide *i* being deployed in monotherapy. If the partner insecticide (*j*) is effective (high dose and/or low levels of resistance) then insecticide *j* effectively kills a large number of mosquitoes. This reduces the number of mosquitoes which were more resistant to insecticide *i*. The selection differential (within generation change in mean resistance) is therefore the difference in the initial mean (dashed black line, *Z̄*_I_) and the mean of the parents (red dashed line, *Z̄_I_*^P^).

### Insecticide Decay and Monotherapy Deployments: Changes in insecticide resistance over a single generation

The implication of insecticide decay when only one insecticide is deployed (monotherapy) is first considered. The parameter ω_τ_^i^ is the killing efficacy of insecticide *i* against a fully susceptible mosquitoes, τ generations after deployment. The value ω_τ_^i^=1 indicates the insecticide is at the recommended dose and kills 100% of susceptible mosquitoes (*Z*_I_ ≤ 0) in a bioassay. Values where ω_τ_^i^>1 indicate the insecticide is above the recommended dose, as may occur with over-spraying for IRS. As insecticides decay there is a reduction in efficacy (ω_τ_^i^) which is calculated using Equations 2d(i) and 2d(ii) in Hobbs & Hastings (2024) and illustrated in Figure 3 presented here.

**Figure 3:**
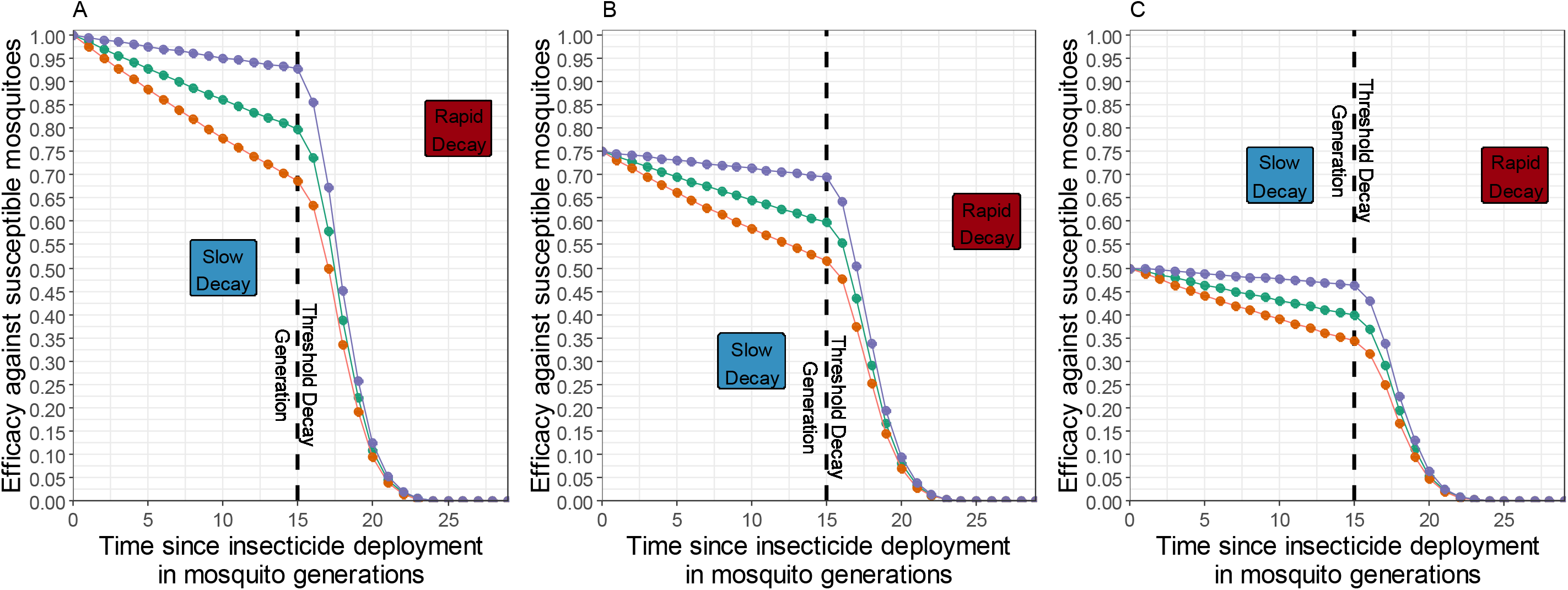
Example Insecticide Decay Profiles for LLINs. As insecticides are deployed over time, there efficacy will decrease as they decay, and insecticide decay rates are likely a function of insecticide chemistry and fabric integrity, giving two-part stages of insecticide decay. The colour of the lines indicates the base decay rate of the insecticide which occurs during the first 15 generations (1.5 years) of deployment and is 0.005 for purple, 0.015 for green (estimated default) and 0.025 for orange, this is the period during which there is slower decay. After 15 generations (1.5 years) (estimated default) the insecticide changes to decaying rapidly, and the decay rate for this period is 0.08 (estimated default). Panel A is when the insecticide is deployed at the recommended dose, as would occur with monotherapy deployments and full-dose mixtures. Panel B and C are when the insecticide is deployed at a reduced dose as may occur with mixture deployments, and the deployed dose is such that the initial efficacy is reduced to either 0.75 (Panel B) or 0.5 (Panel C).

Single generation selection (Figure 1) was conducted across the insecticide efficacy (ω_τ_^i^) space with values from 0 to 1.2 at intervals of 0.1. Using single generation selection does not account for how long insecticides remain at these efficacies which would be dependent on the specific decay profile (Figure 3) but provides a mechanistic understanding of the selection process and the implication of insecticide decay.

The impact of insecticide decay on the response to selection was further assessed across the resistance and exposure space. The initial resistance of the population (measured as bioassay survival, *K_i_^B^*) was set as 0, 5, 10, 20, 50 or 80%. This was run allowing differing levels of exposure (0.2 to 1 at 0.2 intervals), with male and female mosquitoes having the same exposure. Exposure is defined as the proportion of the mosquito population which contact the insecticide(s). Heritability (ℎ^2^) was set at 0.2. Higher and lower heritability values increase and decrease the rate of evolution respectively but does not impact the interpretation of the results (see modified sex-specific Breeder’s equation). Dispersal to/from untreated an untreated refugia (i.e., areas where the insecticide(s) are not deployed) and fitness costs of resistance were not included to simplify interpretation.

All permutations of insecticide efficacy, exposure, and resistance were run (Table 1) allowing us to see how these three parameters interact. The outcome was the response to selection as defined in the Breeder’s equation, i.e., the increase in insecticide resistance over a single mosquito generation.

**Table 1:**
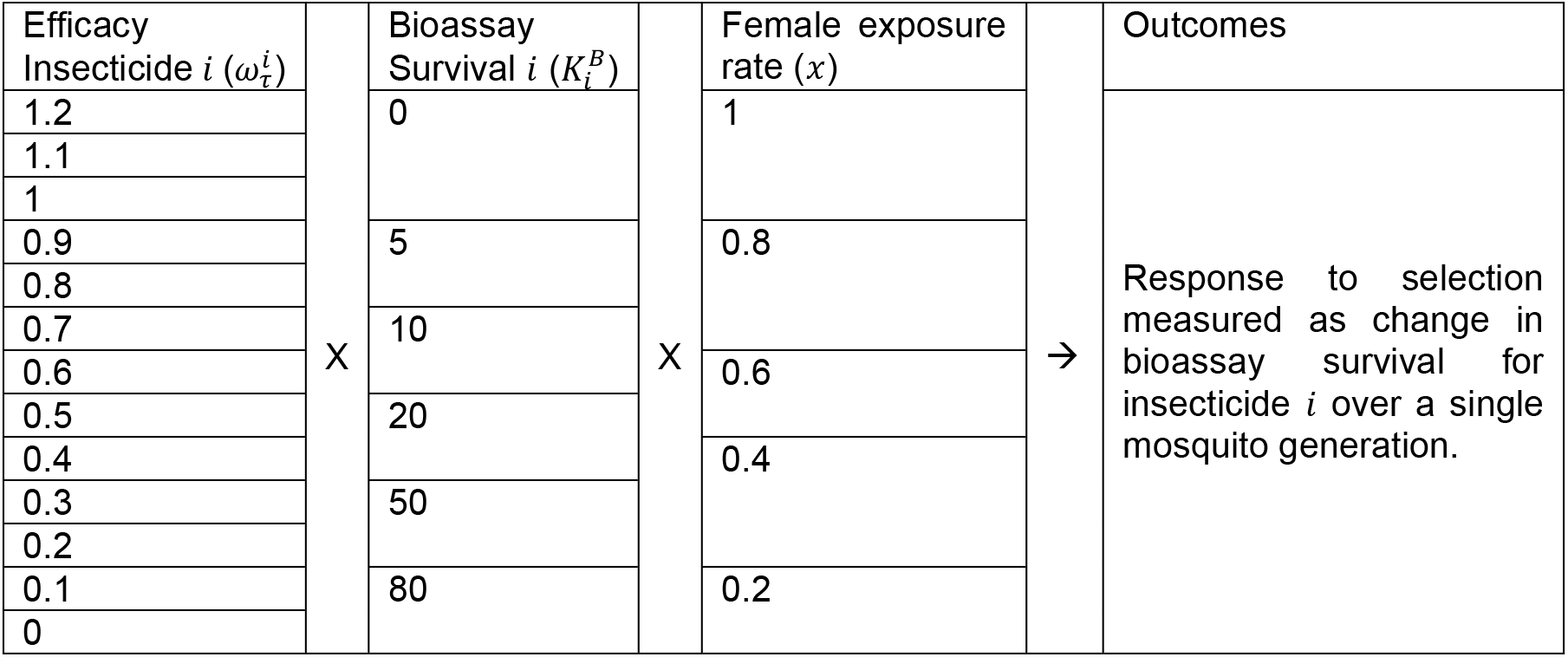
Single generation change in bioassay survival: Monotherapy Deployments.

### Insecticide Decay and Mixture Deployments: change in insecticide resistance over a single generation

The next step evaluates the impact of insecticide decay when two insecticides are deployed in a mixture. Mixtures contain two insecticides (denoted insecticides *i* and *j* in this manuscript) in the same formulation such that mosquitoes are exposed to both insecticides simultaneously. Single generation selection was conducted (Figure 2) across the insecticide efficacy (ω_τ_^i^ and ω_τ_^j^) space. Insecticide *i* and *j* can have different efficacies as would occur if they had different decay profiles or if one was deployed at sub-optimal initial concentrations. Insecticide efficacy parameter space was created at 0.1 intervals ranging from 0-1.2 for each insecticide.

The impact of the exposure (the proportion of the mosquito population which contact the insecticide(s)) and level of resistance to the insecticides was also explored. Exposure to the mixture was set at 0.2 to 1 at 0.2 intervals, with male and female mosquitoes having the same exposure. Initial resistance to the insecticides as bioassay survival (*K_i_*^B^ and *K_j_*^B^) was set as 0, 5, 10, 20, 50 or 80% for each insecticide, allowing the level of resistance to the two insecticides in the mixture to be different. Heritability (ℎ*_I_*^2^ and ℎ*_I_*^2^) to each insecticide was fixed at 0.2. Dispersal to/from refugia and fitness costs were not included to simplify interpretation. All permutations of these values were run (Table 2). This allows us to explore the entirety of the efficacy-resistance-exposure space for mixtures providing detailed mechanistic understanding of the selection process.

**Table 2:** Single generation change in bioassay survival: Mixture Deployments.

|  |  |  |  |  |  |  |  |  |
| --- | --- | --- | --- | --- | --- | --- | --- | --- |
| Efficacy Insecticide $i$<br>($\omega_{\tau}^i$) | | Efficacy Insecticide $j$<br>($\omega_{\tau}^j$) | | Bioassay Survival $i$<br>($K_i^B$) | | Bioassay Survival $j$<br>($K_j^B$) | | Outcomes |
| 1.2 | X | 1.2 | X | 0 | X | 0 | → | • Single generation bioassay change for insecticide $i$.<br>• Single generation bioassay change for insecticide $j$.<br>• Combined total of bioassay change for insecticide $i$ plus bioassay change for insecticide $j$ as measure of the “total amount of selection”. |
| 1.1 |  | 1.1 |  |  |  |  |  |  |
| 1 |  | 1 |  |  |  |  |  |  |
| 0.9 |  | 0.9 |  | 5 |  | 5 |  |  |
| 0.8 |  | 0.8 |  |  |  |  |  |  |
| 0.7 |  | 0.7 |  | 10 |  | 10 |  |  |
| 0.6 |  | 0.6 |  |  |  |  |  |  |
| 0.5 |  | 0.5 |  | 20 |  | 20 |  |  |
| 0.4 |  | 0.4 |  |  |  |  |  |  |
| 0.3 |  | 0.3 |  | 50 |  | 50 |  |  |
| 0.2 |  | 0.2 |  |  |  |  |  |  |
| 0.1 |  | 0.1 |  | 80 |  | 80 |  |  |
| 0 |  | 0 |  |  |  |  |  |  |

The outcome was the response to selection measured as the changes in bioassay survival over a single generation (see Breeder’s Equation and Figure 1). This is measured for both insecticides separately and are added together to get a measure for the total amount of selection. The responses are plotted to visualise regions of the mixture efficacy and resistance space which give higher and lower selection on insecticide *i*, insecticide *j* and both insecticides to gain an understanding of the implication of insecticide decay for mixtures.

### Impact of Insecticide Decay on IRM Strategy Lifespans: multigenerational changes in IR

Clearly an important consideration is how long insecticides remain effective during a deployment, which will depend on the decay profiles of the insecticides (Figure 3). Insecticide decay rates for a standard pyrethroid LLIN collected from the field over time (Toé et al., 2019) were estimated (Supplement 1) to give default parameter values for the decay rate. Insecticide decay for LLINs is likely to be a two-stage process (although the methodology easily allows it to be single stage). During the first stage, the insecticide decays slowly, and after a longevity threshold the second, faster stage of decay rate occurs, for example as the fabric of the LLIN is sufficiently damaged to accelerate the rate of insecticide decay. Many factors can influence the rate of insecticide decay in the field such as the sprayed substrate for IRS (Yewhalaw et al., 2017) and household conditions for LLINs (Mechan et al., 2022). Therefore, we examined a set of illustrative example decay profiles (Figure 3). These decay profiles are used to assess the impact of slower or faster decay rates on the performance of IRM strategies. Simulations were also run where insecticide decay did not occur as this is the usual assumption in most models. Including simulations both with/without decay provides a comparison of models to understand whether inclusion of insecticide decay both quantitatively changed the benefit of IRM strategies, and more importantly whether the inclusion of insecticide decay changed the qualitative conclusion of which strategy performed best.

Simulations were run which allowed the comparison of the two insecticides deployed as monotherapies in sequence, which is generally the default IRM strategy versus the deployment of the same insecticides as a mixture. The properties of the insecticides were the same regardless of the strategy allowing for direct comparisons. The initial dosing for mixtures was set as 100%. The 75% and 50% efficacies were used to emulate the impact of insecticides in mixture being deployed at a reduced dose.

The model additionally allows for the inclusion of cross resistance (*α*_IJ_) as a correlated response (Equation 8b and 8c in Hobbs and Hastings (2024)). The implication of cross resistance between the two insecticides was therefore also explored as this is an important operational concern and is likely to interact with insecticide decay. Previously full-dose mixtures were found to be superior to sequences regardless of the level of cross resistance (Hobbs et al., 2023), however this was without including cross resistance. Cross resistance was included as either positive (*α*_IJ_ = 0.3), negative (*α*_IJ_ = -0.3) or absent (*α*_IJ_ = 0).

For all simulations unless otherwise stated the default parameter values are those detailed in Table 3. Three separate scenarios were conducted to look at the impact of insecticide decay. Scenario 1 assumed both insecticides started at the same resistance level and had the same heritability. This therefore allows the effect of just insecticide decay to be observed. Scenario 2 assumed the insecticides started at different levels of resistance. Scenario 3 assumed the insecticides had different heritabilities. Details of the parameter values used for each scenario are given in Table 3.

**Table 3.**
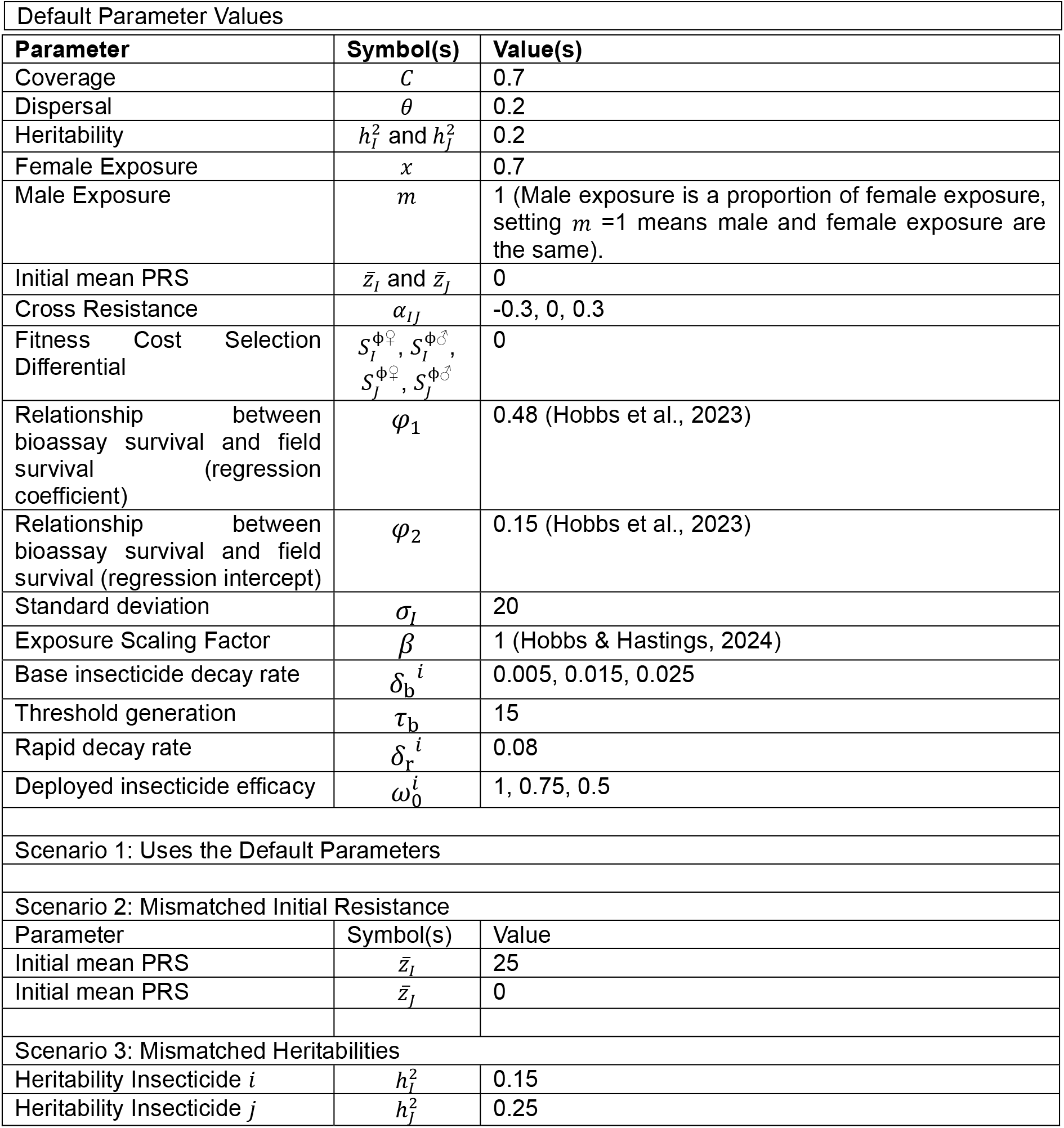
Parameter Values for Simulation Scenarios.

Simulations were run using an insecticide withdrawal threshold of 10% bioassay survival and an insecticide return threshold of 8% bioassay survival (see Table 4 for definitions). Simulations were terminated when (a) they reached the 500-generation maximum duration with one or both insecticides still effective or (b) when no insecticides were available for deployment such that IR to both insecticides remained above the return threshold after withdrawal. For insecticides in mixture, if one of the insecticides reached the withdrawal threshold, the mixture was no longer available to be deployed and the simulation is terminated as the mixture strategy has “failed”. The deployment frequency was 30 generations (∼ 3 years), which is the standard time between LLIN deployments. The outcome was the difference in strategy lifespan (measured in years, assuming 10 mosquito generations per year) between the mixture simulations and the comparator monotherapy sequences simulations.

**Table 4:** Terminology and Definitions.

| Table 4: Terminology and Definitions |  |
| --- | --- |
| Term | Definition |
| Mixture | A single insecticide formulation which contains two insecticides, such that if a mosquito contacts the mixture they inevitably contact both insecticides. |
| Monotherapy | The deployment of a single insecticide. |
| Sequence | Insecticides are deployed until they reach the withdrawal threshold, after which they are withdrawn from deployment and replaced with the next insecticide. |
| Withdrawal Threshold | The level of resistance (measured in bioassay survival), which leads to an insecticide no longer being deployed. In this paper the withdrawal threshold is 10%. |
| Return Threshold | The level of resistance (measured as bioassay survival), which allows a previously withdrawn insecticide to be redeployed. In this paper the return threshold is 8%. |
| Exposure | The proportion of the mosquito population which contact the insecticide(s). |

## Results

### Insecticide Decay and Monotherapy Deployments: change in insecticide resistance level over a single generation

The results first consider the deployment of insecticides as monotherapies, such as happens with the deployment of pyrethroid-only LLINs. Figure 4 shows the impact of reductions in efficacy (e.g., due to insecticide decay) on the resulting singe-generation response to selection across the exposure rate and resistance parameter space. Figure 4 shows that the rate of selection depends on the interaction between exposure, resistance, and insecticide efficacy, such that as the insecticide decays there can be higher levels of selection. We can explain these results mechanistically referring to Figure 1.

**Figure 4:**
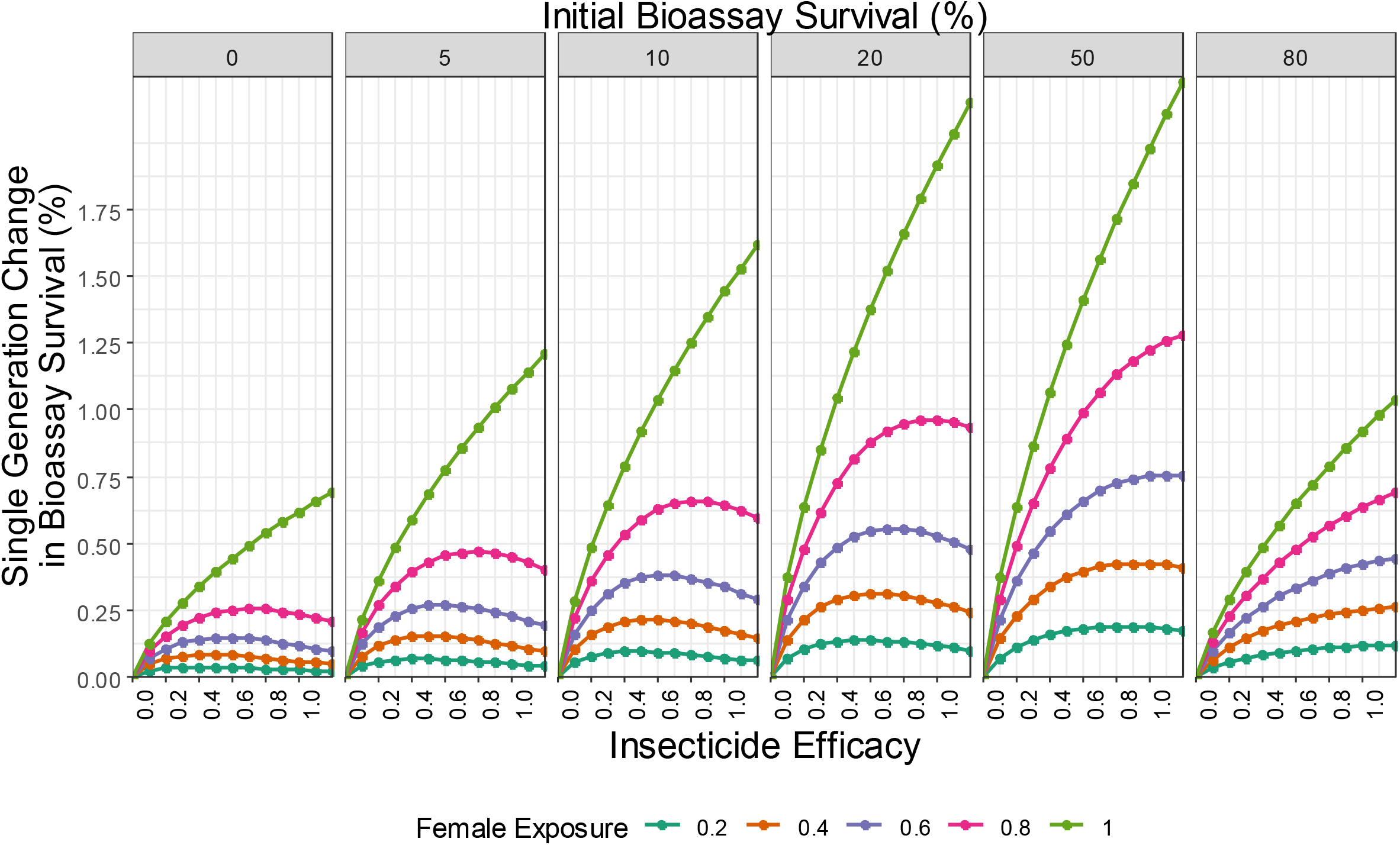
Single generation changes in bioassay survival (%) for monotherapy deployments. Colours indicate the insecticide exposure (where the level of exposure is the same for males and females). The panels are for allowing different levels of initial resistance to the insecticide measured of the population prior to insecticide exposure, measured as bioassay survival.

At high insecticide efficacy (e.g., a newly deployed insecticide, Figure 1: high dose insecticide), the insecticide kills most of the mosquitoes which are exposed to it, and the survivors are only the most resistant individuals in the population. The proportion of the population which avoided the insecticide (and therefore avoiding insecticide selection) makes up most of the final parental population. The resulting response to selection is therefore small. Of course, as female expose increases then so does the response to selection (Figure 4).

When insecticide efficacy is low (e.g., the insecticide has been deployed for a long time with much of the insecticide decaying away, Figure 1: low dose insecticide) most of the mosquitoes exposed to the insecticide survive, with the insecticide killing only the most susceptible individuals. Therefore, the response to selection is small and is further reduced when the final parental population is diluted by mosquitoes which avoided the insecticide.

Between these two extremes, as the insecticide decays, there is a period where the insecticide efficacy means both a significant proportion of the exposed mosquitoes survive and that these mosquitoes are moderately to very highly resistant (Figure 1: Moderate dose insecticide). The proportion of individuals surviving the insecticide is sufficiently large to not be adequately diluted by the mosquitoes which avoided selection. This leads to the response to selection being highest under such conditions of intermediate efficacy (Figure 4), providing some mosquitoes escape selection (if this does not occur, i.e., exposure =1, then the relationship is monotonic; Figure 4).

### Insecticide decay and mixture deployments: change in insecticide resistance level over a single generation

The next step is to consider the impact of insecticide decay for mixtures. We are only considering a single generation so do not need to define a decay profile (e.g., Figure 3) but assume insecticidal decay has reduced the efficacy of one (or both) insecticides and focus on how reduced efficacy drives IR. Figure 5 is a heat map of the responses to selection for mosquitoes exposed to a mixture of both insecticide *i* and insecticide *j* (assuming, for illustration, that 60% of mosquitoes in the population are exposed to the insecticide across the efficacy and resistance space). Figure 5 contains 3 panels, one for the response to each insecticide separately and a third for the total response (indicating the total amount of selection on the population). Plots for additional exposure rates are found in Supplement 2 (Figure S2.1) although their qualitative interpretation is similar as for Figure 5, albeit higher exposures have larger single generation changes and lower exposures have smaller single generation changes. When exposure is 1 no mosquitoes escape exposure and therefore increasing the efficacy always increases the amount of selection (Supplement 2, Figure S2.2).

**Figure 5:**
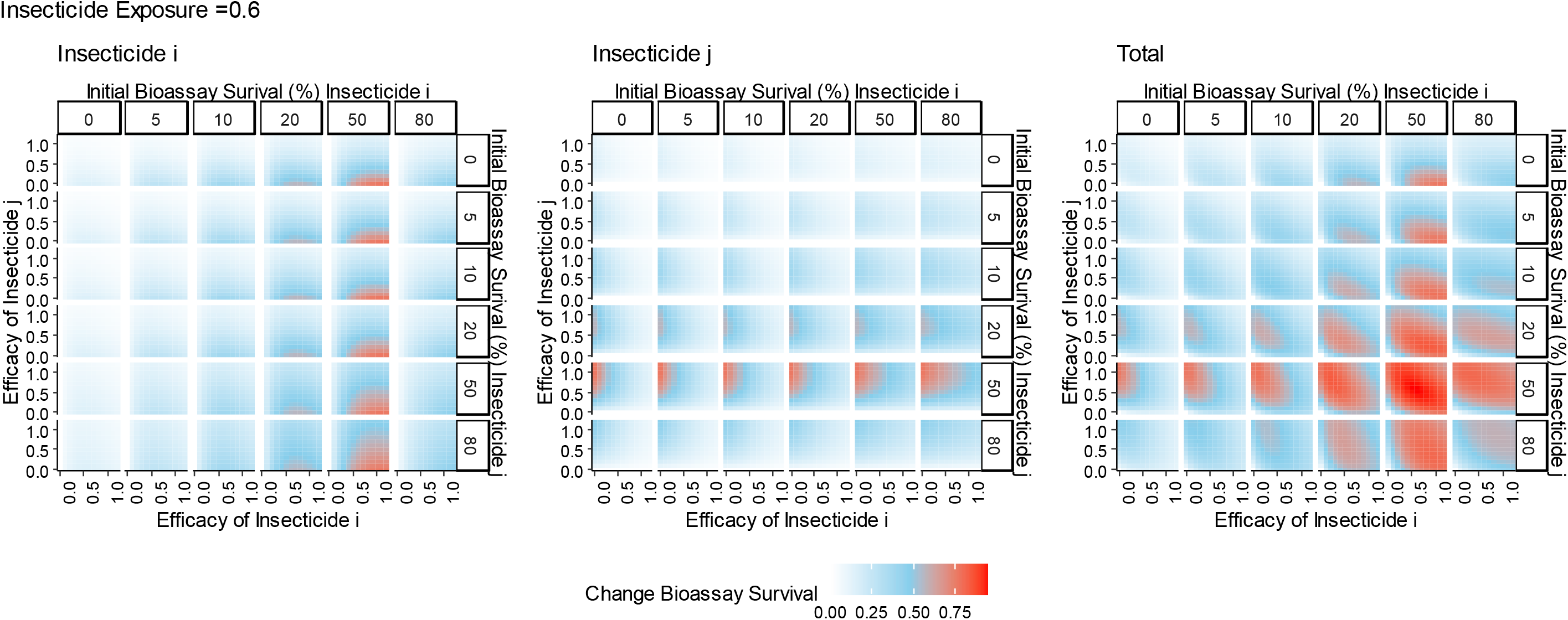
Single generation changes in bioassay survival (%) for mixture deployments with exposure = 0.6. Interpretation: Red values indicate larger changes in bioassy survival so are worse for IRM. Pale blue values indicate smaller change in bioassay survival and so are better for IRM. Left plot: Change in bioassay survival for insecticide *i* only. Middle plot: Change in bioassay survival for insecticide *j* only. Right plot: Total combined change in bioassay survival (change bioassay survival insecticide *i* + change bioassay survival insecticide *j*). The x axis is the efficacy of insecticide *i* and y axis is the efficacy of insecticide *j*. Panels (left-right): the amount of initial resistance to insecticide *i* as measured in a bioassay. Panels (top-bottom): the amount of initial resistance to insecticide *j* as measured in a bioassay.

Figure 5 shows that the response to selection is highest where there is already resistance to both insecticides and/or (equivalently) the efficacy of both insecticides is reduced. This can be mechanistically explained using Figure 2. In Figure 2, the mortality induced by insecticide *i* remains constant between each panel, with the additional mortality induced by insecticide *j* changing. It shows that additional mortality imposed by insecticide *j* reduces the response to selection for insecticide *i*. This is because insecticide *j* kills additional mosquitoes in the exposed group and therefore increases the proportion of the final parental population which are from the unexposed group. Of course, there would be simultaneously selection for resistance to insecticide *j* also.

Of immediate policy concern is the implication for next-generation mixture LLINs. With next-generation mixture LLINs, a novel insecticide (with little to no resistance to it) is deployed with a pyrethroid insecticide (where there is likely to already be substantial levels of resistance). Figure 6 focuses on next-generation mixture LLINs and looks at how the rate of selection on the novel (insecticide *i*) insecticide is impacted by being in mixture with differing levels of resistance to the pyrethroid (insecticide *j*). For simplicity of interpretation selection rates were split into quartiles (“low”, “moderate”, “high”, “very high”), and was done to highlight the regions of higher selection which increase as the resistance to the pyrethroid (insecticide *j*) partner increases. It can be clearly seen that mixture efficacy space area of the “low selection rate” reduces as resistance to the pyrethroid (insecticide *j*) increases. From both Figure 5 and 6, it should be highlighted that selection appears to be highest when efficacy is at 0.5, which may be the efficacy of half-dose mixtures.

**Figure 6:**
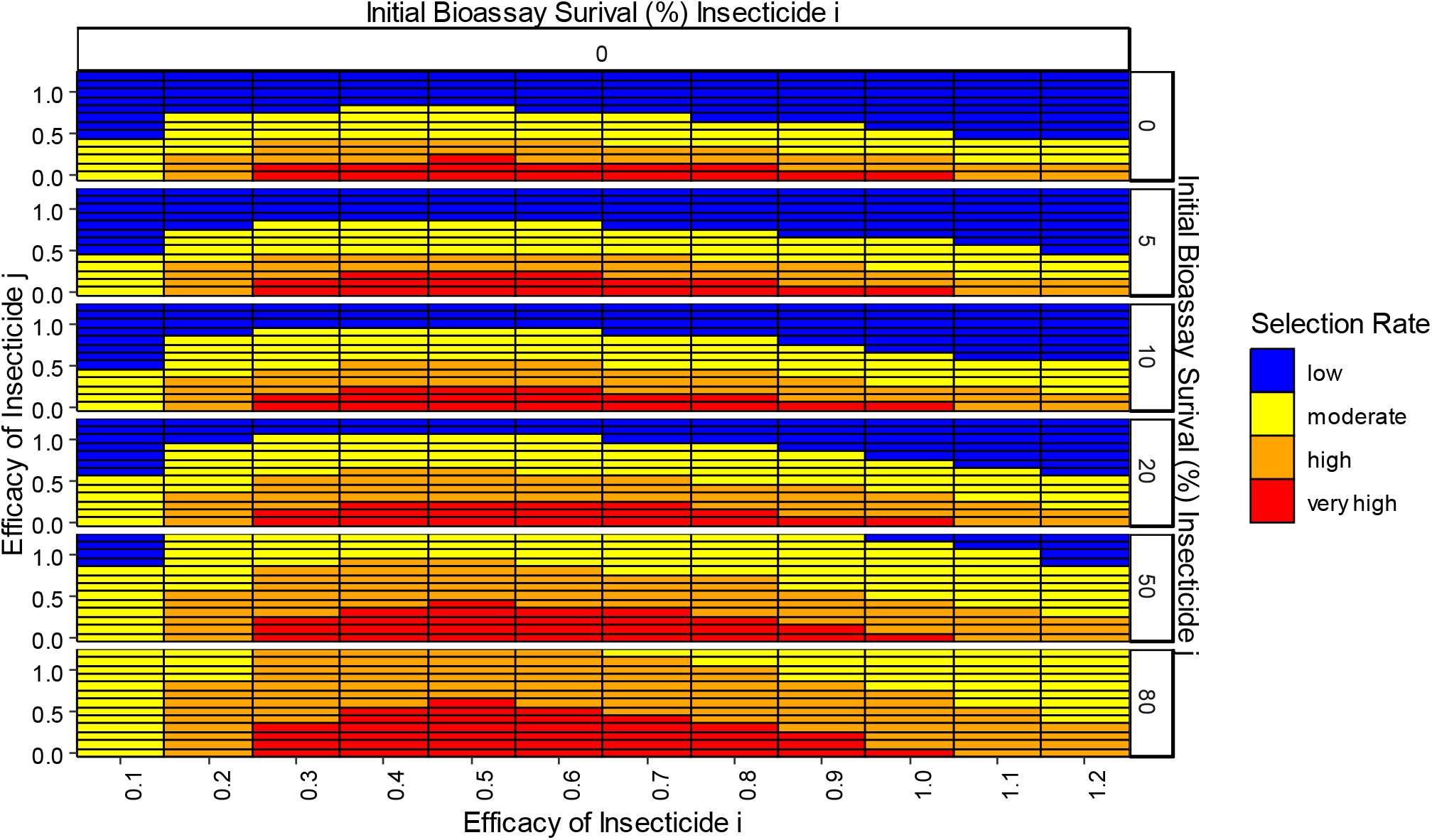
A simplified schema of the rate of selection on one insecticide (insecticide *i* in this case) as a consequence of increasing resistance to the partner insecticide when deploying mixtures with exposure = 0.6. The changes in bioassay survival were measured over the whole parameter space shown in the figure, then classed into quartiles: “low” (blue quartile), “moderate” (yellow quartile “high” (orange quartile) and “very high” (red quartile). The descriptors and bins are essentially arbitrary and are used to consider the implication of insecticide decay and what situations should generally be encouraged or avoided. This figure easily demonstrates the qualitative implication of insecticide decay i.e., that selection for resistance to insecticide *i* is highest when (1) efficacy of insecticide *i* is intermediate (lower x axis) and (2) efficacy of its partner, insecticide *j* is low either because it is inherently less effective (left y axis) or because substantial resistance is already present (right y axis).

### Impact of Insecticide Decay on IRM Strategy Lifespans: multigenerational changes in IR

Simulations were run to investigate the long-term impact of insecticide decay on the performance of mixtures versus monotherapy sequences over realistic timescales (up to 500 mosquito generations, equivalent to 50 years).

Scenario 1 (Table 3; Figure 7) includes insecticide decay alone, without the additional complexities of mis-matched initial resistance or mis-matched heritability. Failure to include insecticide decay appears to overestimate the effectiveness of full-dose mixtures for IRM when compared to monotherapy sequences, albeit still outperforming the monotherapy sequence strategy. At higher rates of decay, the benefit of full-dose mixtures became smaller. The reduced dose mixtures performed poorly regardless of whether insecticide decay was included or not.

**Figure 7:**
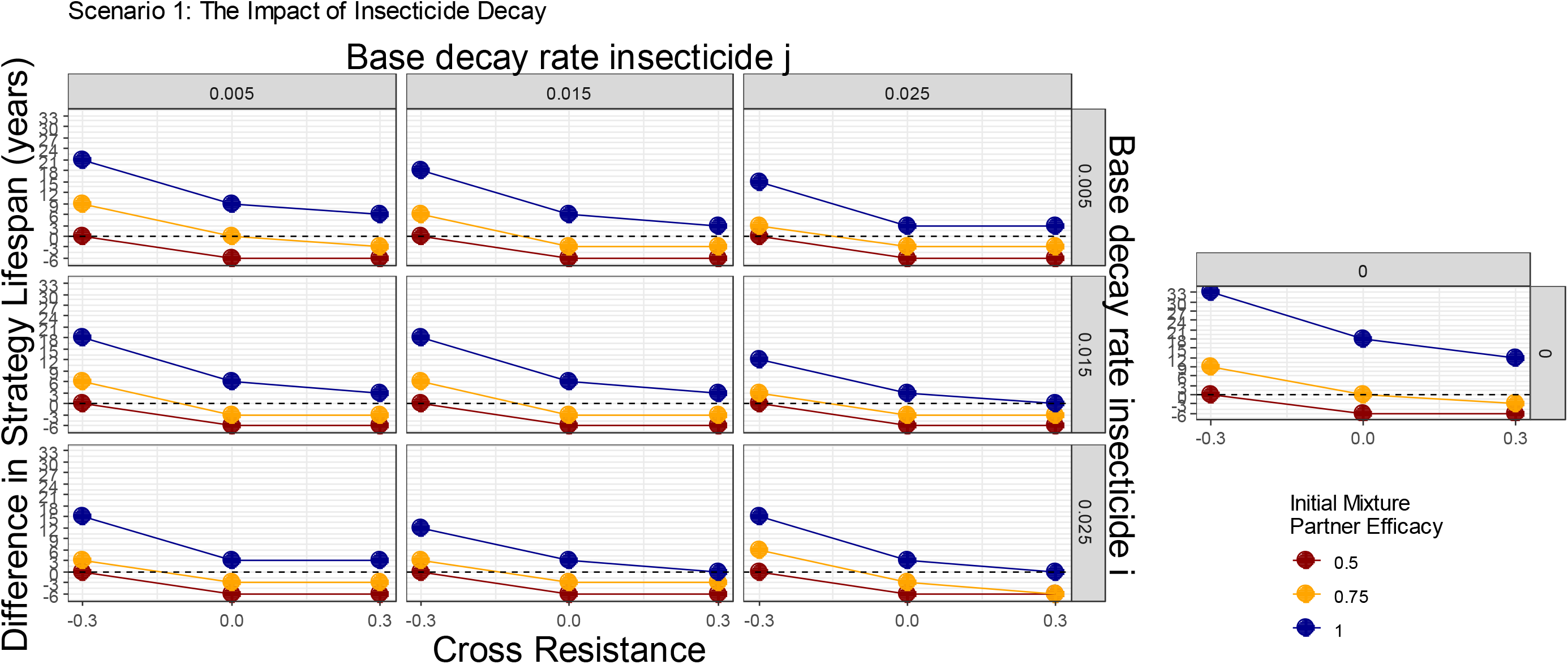
Scenario 1 – Impact of including insecticide decay in simulations evaluating mixtures versus monotherapy sequences. The colours indicate the deployed efficacy of each insecticide when in mixture. The dashed line indicates the mixture strategy and monotherapy sequence strategy had the same strategy lifespans. Values above this line indicate the mixture strategy had a longer strategy lifespan and values below this line indicates the monotherapy sequence strategy had the longer strategy lifespan. The panels top-bottom are the base decay rate for insecticide *i*, and the panels left-right are the base decay rate for insecticide *j*. The x axis (bottom) is the degree of cross resistance between the two insecticides. The inset graph is the simulation where insecticide decay does not occur and has the same axis labels.

Scenario 2 (Table 3; Figure 8) extends Scenario 1 to allow two insecticides with mis-matched initial resistance. Concerningly, when there was some pre-existing resistance to one of the mixture partners there are decay profiles where the full-dose mixture performs worse than monotherapy sequences. Scenario 3 (Table 3; Figure 9) extends Scenario 1 by allowing mis-matched heritabilities for the resistance traits. Here we see that there are decay profiles where again full-dose mixtures perform worse than monotherapy sequences.

**Figure 8:**
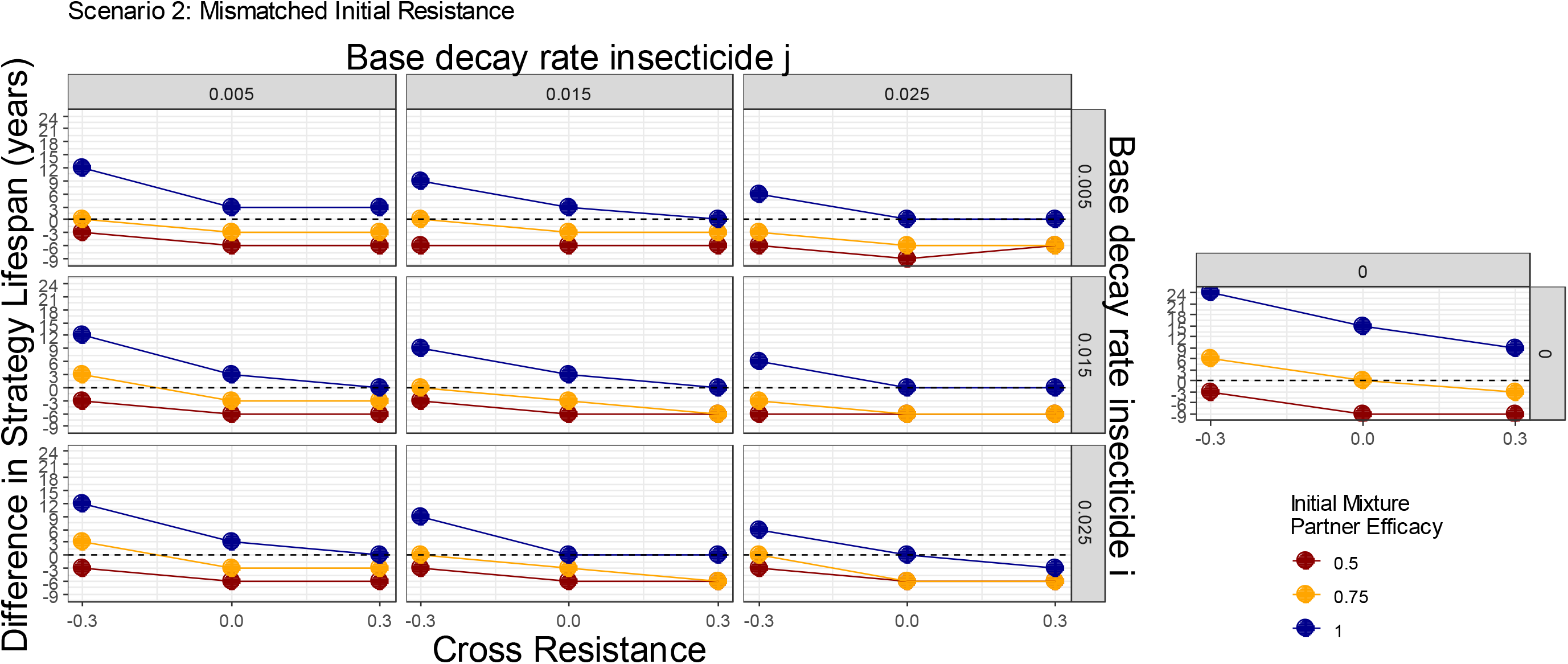
Scenario 2 – Impact of including insecticide decay in simulations evaluating mixtures versus monotherapy sequences with mismatched initial resistances. The colours indicate the deployed efficacy of each insecticide when in mixture. The horizontal dashed line indicates a difference in lifespan of zero i.e. the mixture strategy and monotherapy sequence strategy had the same strategy lifespans. Values above this line indicate the mixture strategy had a longer strategy lifespan and values below this line indicates the monotherapy sequence strategy had the longer strategy lifespan. The panels top-bottom are the base decay rate for insecticide *i*, and the panels left-right are the base decay rate for insecticide *j*. The x axis (bottom) is the degree of cross resistance between the two insecticides. The inset graph is the simulation where insecticide decay does not occur and has the same axis labels.

**Figure 9:**
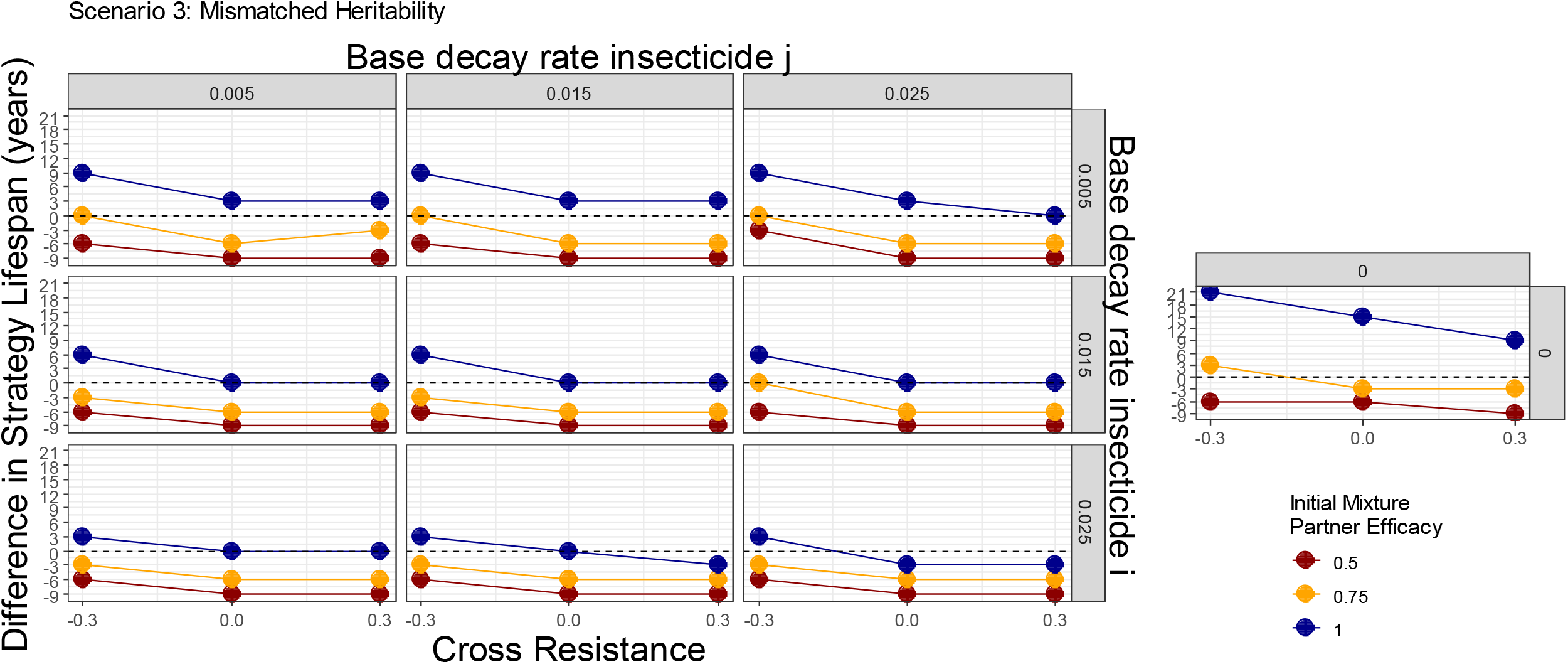
Scenario 3 – Impact of including insecticide decay in simulations evaluating mixtures versus monotherapy sequences with mismatched heritabilities. In this scenario the two insecticides had different heritabilities. The heritability for insecticide *i* (*h_I_*^2^) was 0.15 and the heritability for insecticide *j* (*h_J_*^2^) was: 0.25. The colours indicate the deployed efficacy of each insecticide when in mixture. The dashed line indicates the mixture strategy and monotherapy sequence strategy had the same strategy lifespans. Values above this line indicate the mixture strategy had a longer strategy lifespan and values below this line indicates the monotherapy sequence strategy had the longer strategy lifespan. The panels top-bottom are the base decay rate for insecticide *i*, and the panels left-right are the base decay rate for insecticide *j*. The x axis (bottom) is the degree of cross resistance between the two insecticides. The inset graph is the simulation where insecticide decay does not occur and has the same axis labels.

The generalities across these three scenarios are that (a) failure to include insecticide decay consistently overestimates the benefit of full-dose mixtures (b) reducing the dose each partner in a mixture is additionally concerning (c) combining these two factors is especially concerning.

## Discussion

Insecticide decay is frequently highlighted as an important consideration in the deployment of insecticides in public health, especially with regards to mixtures (Rex Consortium, 2013). Yet insecticide decay has been missing from models (South et al., 2020), a limitations often noted when investigating mixtures e.g. (Curtis & Hill, 1993; Roush, 1989). A dynamic model of polygenic insecticide selection (Hobbs & Hastings, 2024) was used, which allowed the impact of insecticide decay to be explored under both single- and multi-generation selection scenarios using both monotherapy and mixture deployments.

### Impact of Insecticide Decay on Monotherapies

Monotherapy deployments (such as using pyrethroid only LLINs) can include periods post-deployment where the overall selection is higher than when the insecticide was newly deployed (Figure 4), and the durations of these periods depend on the decay profile of the insecticide (Figure 3). A similar result was found when modelling monogenic systems (South et al., 2020).

### Impact of Insecticide Decay on Mixtures

Issues concerning insecticide decay and mixtures have long been highlighted as an issue (Rex Consortium, 2013) despite never being explicitly evaluated. Figure 5 shows how the rate of selection for resistance to one insecticide in a mixture is modulated by the efficacy and resistance status of the partner insecticide. Perhaps the most concerning result is that the rate of the selection (to the mixture as a whole) appears to be fastest when both insecticides are at lower efficacies (approximately 0.5); worryingly this may be the approximate efficacy of half-dose mixtures. The decay of mixtures in the field y will of course determine how long the mosquitoes are under these periods where selection is higher or lower; and worryingly these reductions in insecticide efficacy are likely to coincide with reductions in disease control.

### Impact of Insecticide Decay on IRM Strategy Performance

Insecticide decay occurs regardless of the IRM strategy used. However, the important question is whether the inclusion of insecticide decay in simulations change the choice of which IRM strategy should be deployed? Our results show that this is an important issue for impacting the rate of selection (Figures 4 and 5) but is also important in determining whether a strategy is effective relative to another (Figure 7, 8 and 9). The failure to include insecticide decay in previous models evaluating mixtures versus monotherapies (e.g., Curtis, 1985; Hobbs et al., 2023; Madgwick & Kanitz, 2022b; Mani, 1985; South & Hastings, 2018) may unfortunately have be giving overly optimistic estimates of the performance of full-dose mixtures for IRM. This was, of course, recognised and noted by these authors but, to our knowledge, the work presented here is the first to include the effect.

### Impact on Personal Protection and Disease Transmission

The inclusion of an insecticide decay parameter in transmission models is common (e.g., Briët et al., 2013; Churcher et al., 2016; Yakob et al., 2011) to account for the expectation of reduced personal protection and impact on disease transmission over time. However, at the same time, these decaying insecticides are likely still providing selection for insecticide resistance further compounding reductions in personal protection and disease transmission, an effect that transmission models generally do not include. These low levels of insecticide efficacies can lead to increases in insecticide resistance over successive generations, potentially compounding the issue of reduced insecticide efficacy on personal protection and disease transmission. An interim IRM strategy may be to increase the rate at which worn-out standard pyrethroid-only LLINs are replaced with new ones. Damaged nets can be checked through visual observation (WHO, 2013), to identify LLINs where there is no longer sufficient insecticide is more challenging.

### Caveat: Degree of Control

These mechanistic descriptions of how mixtures (Figure 2) work also help demonstrate the additional kill concept (Madgwick & Kanitz, 2022a), e.g., the orange parts of Figure 2. Mixtures (especially when both insecticides are at high dose) would be expected to kill more individuals than monotherapies and this benefit of increased population suppression is not usually considered in the evaluation of IRM strategies (Madgwick & Kanitz, 2022a), especially those which evaluate public health interventions as linking resistance and transmission is a complex challenge (Barbosa et al., 2018). The orange sections of Figure 2 demonstrate the additional kill provided by a partner insecticide and shows additionally how this decreases the population size of mosquitoes (likely providing an increased vector control benefit and subsequently control of transmission), and also increasing the relative contribution of mosquitoes which were not selected as they avoided the insecticides (the IRM benefit).

## Conclusion

Insecticide effectiveness and insecticide decay are key considerations when evaluating IRM strategies. Despite being frequently highlighted as an issue, the inclusion of insecticide decay in models has often been absent. As insecticides decay, there can be changes in the rate of selection for resistance. The absence of insecticide decay from previous models appears to have been providing overly optimistic estimates of the performance of full-dose mixtures versus monotherapy deployments.

## Acknowledgements

We would like to acknowledge the advice of David Weetman for help grounding the model in biological and operational relevance. We would also like to acknowledge the Vector Informatics and Genomics group at the Liverpool School of Tropical Medicine for their general critiques of the model methodology.

