## Supplement 1 for "The impact of insecticide decay on the rate of insecticide resistance evolution for monotherapies and mixtures"

### Supplement 1: Estimation of Insecticide Decay Rates for LLINs:

For the insecticide decay we provide values for a default decay profile for an LLIN insecticide and an IRS insecticide. We make note that insecticide decay is likely to be highly heterogenous between different locations due to climatic conditions. But also note that local level conditions will also play a role in the decay rates of insecticides. This is perhaps most notable for IRS where the substrate the insecticide is sprayed on has a large effect on how fast or slow the efficacy of the insecticide declines. Our preference then is rather than attempting to overfit decay profiles to specific insecticides under specific conditions we instead construct default decay profiles, which describe how insecticide efficacy decays over time. And then use the parameter values for default as a baseline to which we can adjust and ask what is the impact of the insecticide decaying faster/slower and or rapidly decaying sooner/late? From these more general global conclusions can be assessed. Using data obtained from Toé et al (2019), which had field collected LLIN assessed in wire ball assays against a fully susceptible pyrethroid susceptible strain (Kisumu; Bioassay Survival = 0%). LLIN collections from the field were made at 0, 6, 12, 18 and 24 months (Toé et al., 2019). Which in our mosquito generation scale correspond to 0, 5, 10, 15 and 20 mosquito generations respectively (we assume 10 mosquito generations per year). As initial wire ball bioassay survival was ~10% we converted the initial starting insecticide efficacy to 0.9. Equation 1b(ii) from Hobbs & Hastings (2024) was used to estimate the base decay rate, rapid decay rate and threshold generation. A broadly matching decay profile was achieved with base decay = 0.015, rapid decay = 0.08 and threshold generations = 15 (Figure S1.1). These values are therefore the default decay rates and decay thresholds, around which can vary. We can therefore vary these values by either increasing or decreasing to assess how the speed (and shape) of the decay profile impacts the rate of insecticide resistance evolution, and what the implications are of insecticides which decay faster and/or slower.

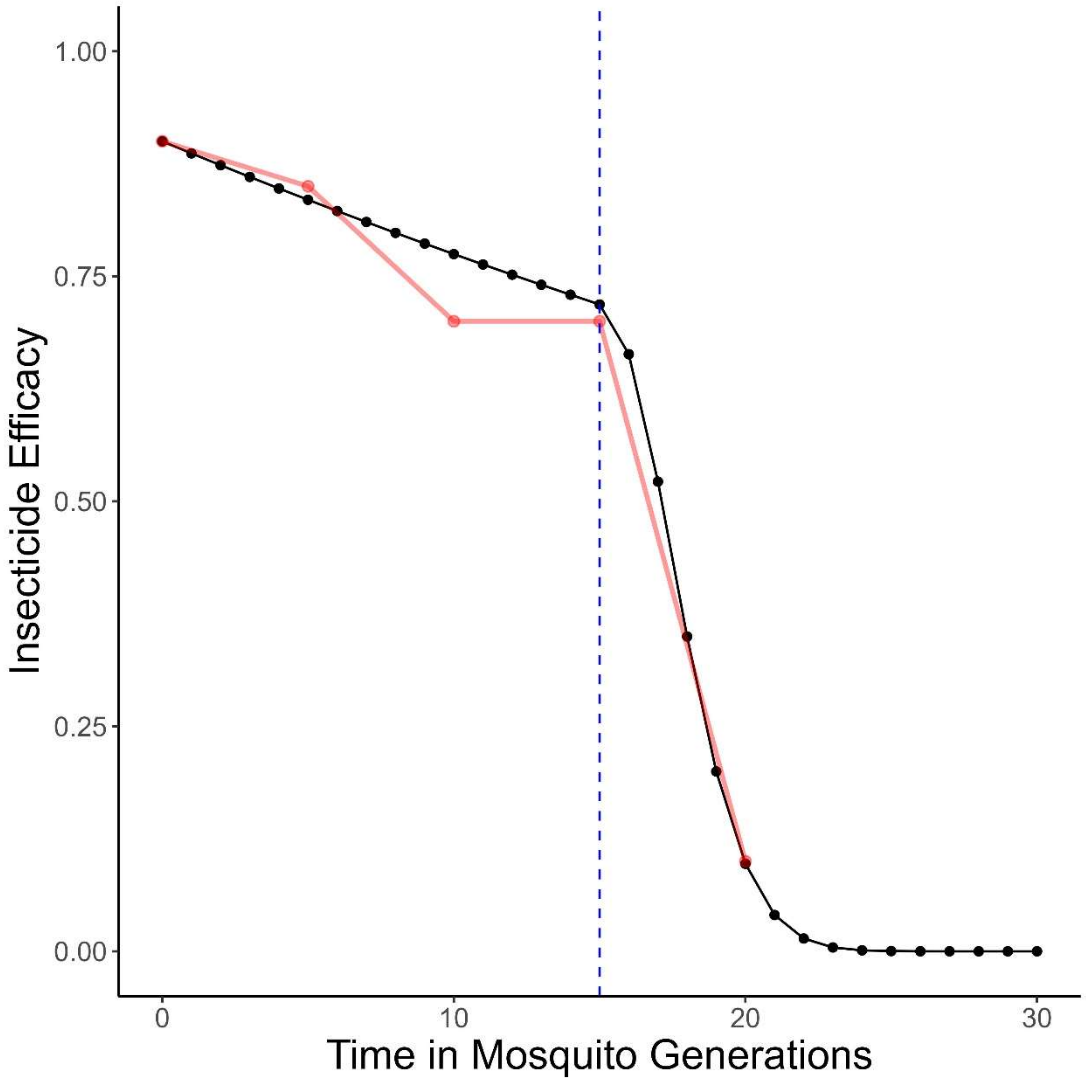


**Figure S1.1: Estimating Insecticide Decay Rates.** Red dots and line is the measured insecticide efficacy from wire-ball assays from field collected LLINs over time. Black line is calculated insecticide efficacy with a base decay rate of 0.015, a rapid decay rate of 0.08 and a threshold generation of 15 (~1.5 years).
