## Supplement 2 for "The impact of insecticide decay on the rate of insecticide resistance evolution for monotherapies and mixtures"

### Supplement 2: Sensitivity Analysis of Mixture Efficacy and Insecticide Exposure

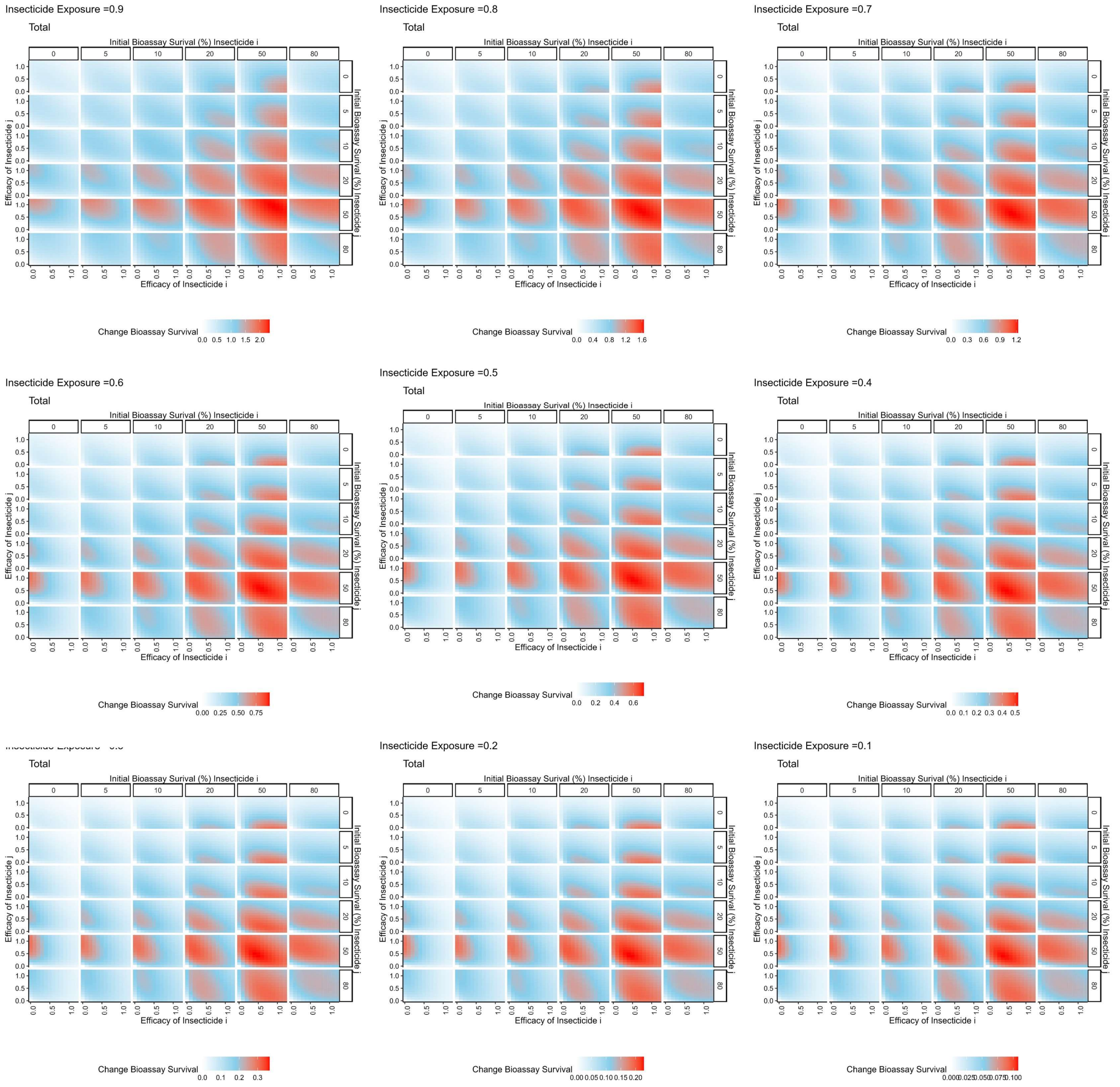


**Figure S2.1: Single generation changes in bioassay survival (%) for mixture deployments with different exposure.** Interpretation: Red values indicate larger changes in bioassy survival so are worse for IRM. Pale blue values indicate smaller change in bioassay survival and so are better for IRM. Plots show the total combined change in bioassay survival (change bioassay survival insecticide $i$ + change bioassay survival insecticide $j$). The x axis is the efficacy of insecticide $i$ and y axis is the efficacy of insecticide $j$. Within each plot: Panels (left-right): the amount of initial resistance to insecticide $i$ as measured in a bioassay. Panels (top-bottom): the amount of initial resistance to insecticide $j$ as measured in a bioassay.

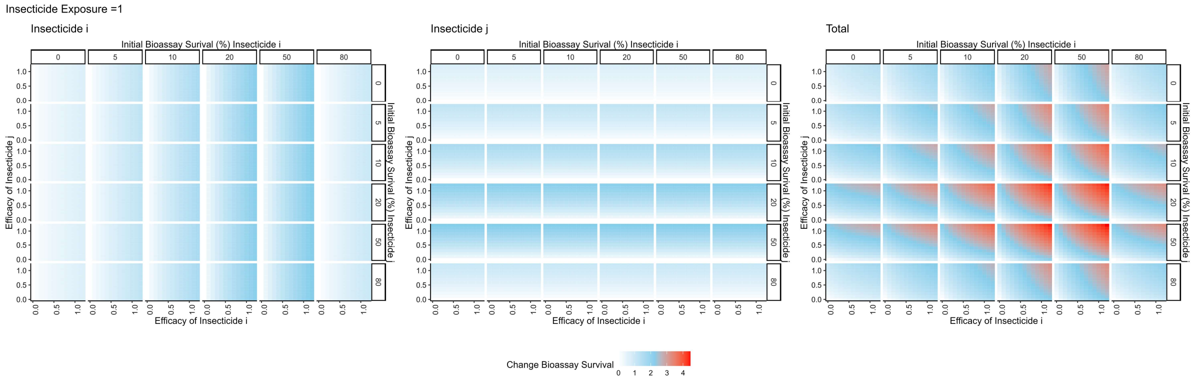


**Figure S2.2: Single generation changes in bioassay survival (%) for mixture deployments with exposure = 1.** Interpretation: Red values indicate larger changes in bioassy survival so are worse for IRM. Pale blue values indicate smaller change in bioassay survival and so are better for IRM. Left plot: Change in bioassay survival for insecticide $i$ only. Middle plot: Change in bioassay survival for insecticide $j$ only. Right plot: Total combined change in bioassay survival (change bioassay survival insecticide $i$ + change bioassay survival insecticide $j$). The x axis is the efficacy of insecticide $i$ and y axis is the efficacy of insecticide $j$. Panels (left-right): the amount of initial resistance to insecticide $i$ as measured in a bioassay. Panels (top-bottom): the amount of initial resistance to insecticide $j$ as measured in a bioassay.
